# Ecological synchrony in human-modified landscapes under a changing climate

**DOI:** 10.1101/2024.11.09.621137

**Authors:** Yiluan Song, Mallory Barnes, Dawn M. Browning, K. Ann Bybee-Finley, Kyla M. Dahlin, Stephan B. Munch, Guillermo E. Ponce-Campos, Casey Youngflesh, Benjamin Zuckerberg, Kai Zhu

## Abstract

Different aspects of ecological systems, biotic or abiotic, often fluctuate in coordinated patterns over space and time. Such high concordance between ecological processes is often referred to as ecological synchrony. Anthropogenic activities, including and beyond climate change, have the potential to alter ecological synchrony by disrupting or enhancing existing synchrony. Despite many local studies, we have a limited systematic understanding of how ecological synchrony is shaped by management in human-dominated landscapes at regional to continental scales. From a macrosystems perspective, we review how anthropogenic activities, particularly beyond climate change, alter ecological synchrony across levels of ecological organization, from the ecosystem level to the population level. For each level, we use a large-scale case study to demonstrate ways to quantify the impacts of human modifications on synchrony using big data from remote sensing, surveys, and observatory networks. For example, we detected possible homogenization of population dynamics of bird species in North America. These changes in ecological synchrony, although in different forms, often represent challenges to ecological and social systems. Collaborative research efforts that integrate emerging open data streams moving forward will be able to provide insights into the effects of different anthropogenic drivers and the consequences of changes in synchrony.

## 1. Changing ecological synchrony in the Anthropocene

Incorporating the temporal dimension of ecological changes has become increasingly critical to address in global change studies at various scales. For example, landscape ecologists have long recognized the role of metapopulation dynamics, in addition to population size, in determining the persistence of a species (A. Liebhold et al., 2004a). Community ecologists have revealed the importance of complementarity over time in addition to species richness in maintaining community stability (Blüthgen et al., 2016; Valencia et al., 2020). Ecosystem ecologists are equipped to examine the coordination between ecosystem processes synthesizing time series in ecology, biogeochemistry, and hydrology (Seybold et al., 2021). These studies that address ecological changes on the temporal dimension, can be synthesized across scales under the emerging study area of ecological synchrony (Fig. 1).

**Figure 1.**
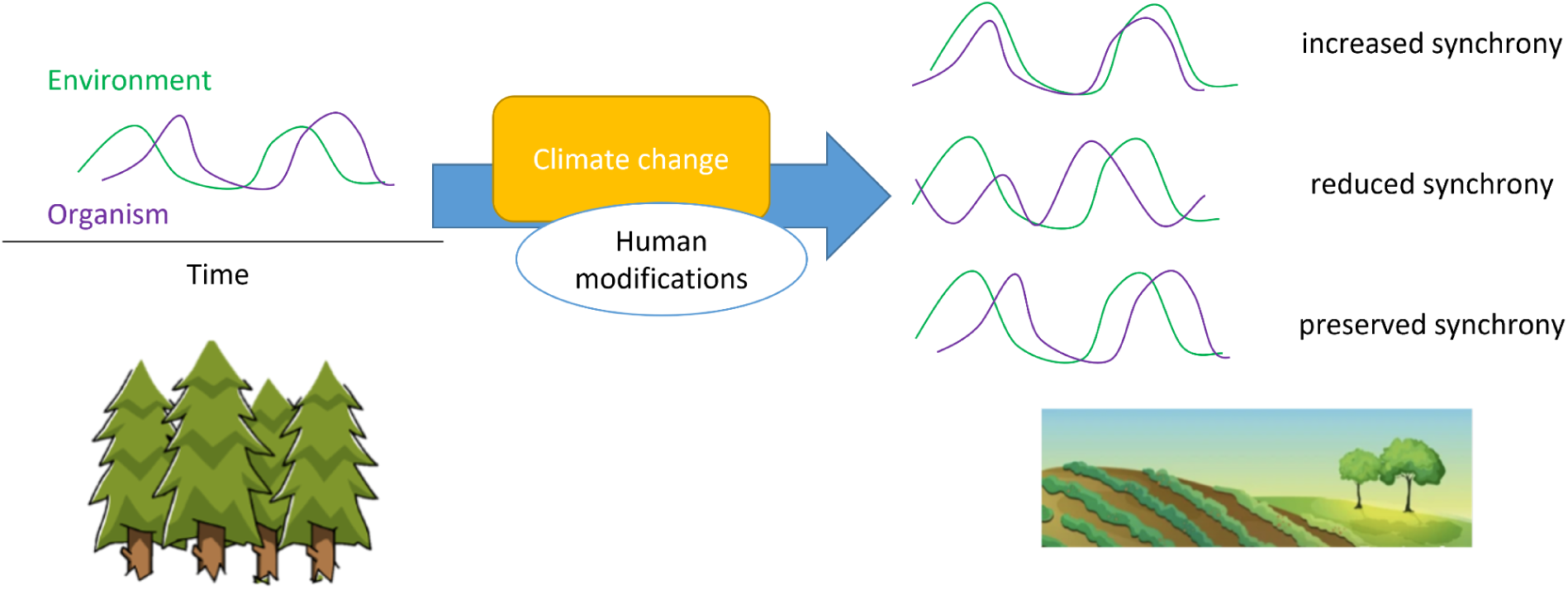
Schematic representation of various possible changes in ecological synchrony with climate change and human modifications.

Ecological synchrony (hereafter referred to as “synchrony”) can be thought of as how different aspects of ecological systems fluctuate in coordinated patterns over space and time. Importantly, synchrony can be found across scales from a macrosystems perspective (Fig. 2), referring to individual behavior, population fluctuations within (Saino et al., 2011; Visser et al., 2012) or between species (Rafferty et al., 2015; Renner & Zohner, 2018), and interactions between abiotic and biotic components (Beard et al., 2019). Synchrony might be found between different systems at the same location, such as prey and predators (Vasseur & Fox, 2009), herbivores and plants (van Asch & Visser, 2007), organisms and abiotic environments (Özkan et al., 2016) and many more. Synchrony might also take place at disjunct locations across space (i.e., spatial synchrony) (A. Liebhold et al., 2004a), including the masting behavior of plants (Koenig et al., 2015; Lamontagne & Boutin, 2007), regional insect outbreaks (A. M. Liebhold et al., 2012) and wildlife disease outbreaks (Princé et al., 2018). Synchrony (and sometimes asynchrony, Box 1) plays an important role across multiple scales of ecology, ranging from population persistence (K. C. Abbott, 2011) to species interactions (Kharouba et al., 2018) to ecosystem functioning (Wang et al., 2021).

**Figure 2.**
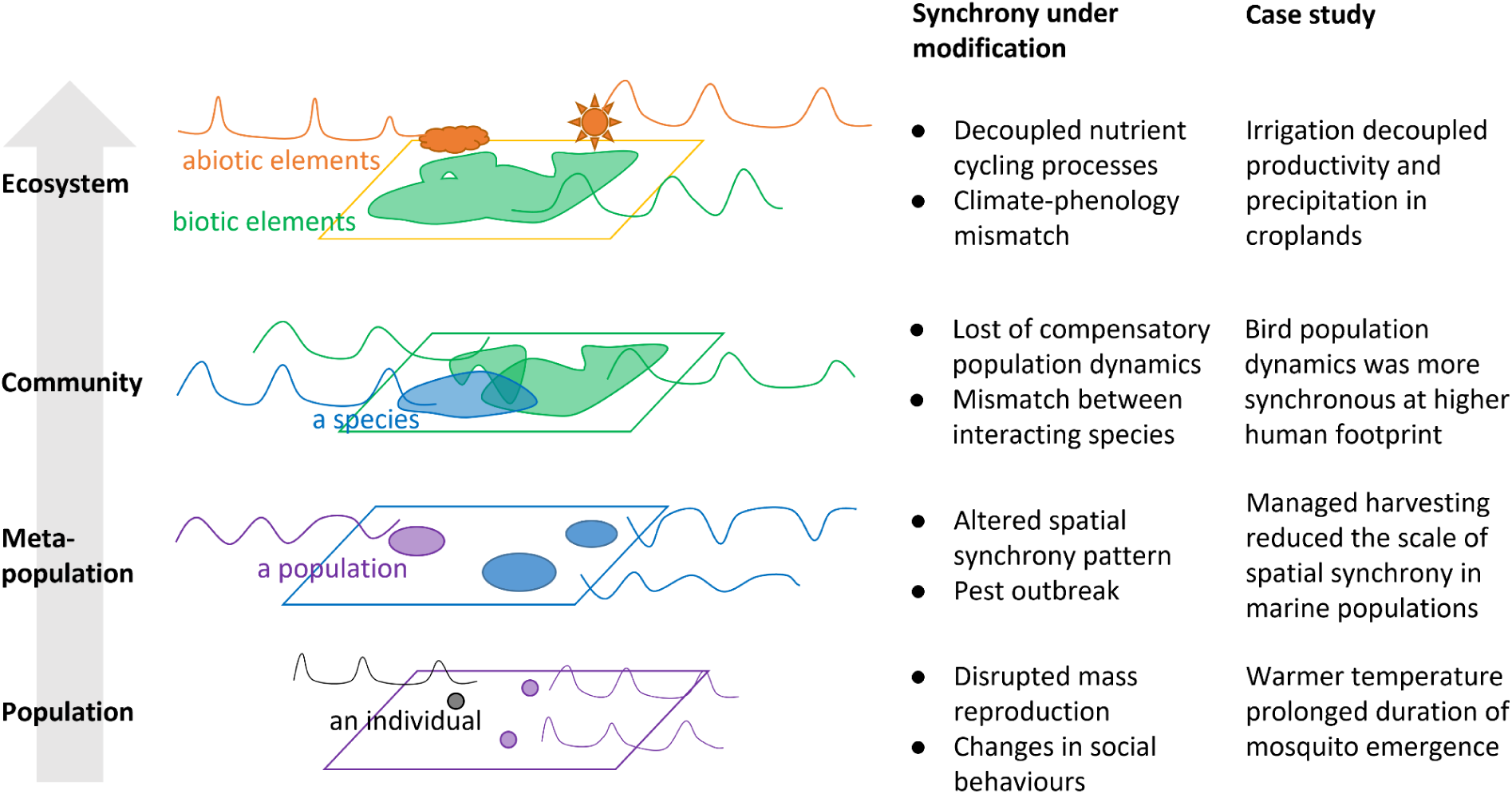
Human activities potentially alter ecological synchrony across levels of ecological organization. Examples from literature (section 2) and case studies (section 3) for each level are provided.

Anthropogenic climate change has changed the pace of many aspects of ecological systems, such as temperature fluctuations, phenology, and population cycles (Post, 2013). Furthermore, climate change can alter synchrony through diverse mechanisms operating across various ecological, spatial, and temporal scales. On the one hand, climate change has been found to disrupt ecological synchrony, as the types and thresholds of environmental triggers and the sensitivity of organisms to these triggers vary (Youngflesh, Socolar, et al., 2021). For example, disparate phenological changes between species in response to changing climate (Kharouba et al., 2018) can shift the outcome of their interactions, resulting in reduced fitness or ecosystem services (Cushing, 1969; Kharouba & Wolkovich, 2020). On the other hand, climate change can enhance the synchrony of a wide range of ecological dynamics at a single location (Youngflesh, Li, et al., 2021) or across multiple locations (Koenig & Liebhold, 2016), often acting through an underlying abiotic driver such as extreme weather (Rahmstorf & Coumou, 2011).

The implications of changes in synchrony for the ecosystem are far-reaching. The loss of synchrony between species and favorable (abiotic or biotic) environmental conditions such as resource availability may negatively affect their persistence (Miller-Rushing et al., 2010; Visser & Gienapp, 2019), but synchronized population dynamics can also increase the risk of extinction during extreme events (Heino et al., 1997; Post & Forchhammer, 2004), undermine demographic rescue among interacting subpopulations (Koenig & Liebhold, 2016), or promote more significant insect outbreaks (Sheppard et al., 2016).

The role of human disturbance and management is often lacking in discussions about ecological synchrony. In addition to anthropogenic climate change, human activities, such as land-use change or habitat management, modify ecological synchrony in several ways (Fig. 1). Management practices can either promote, reduce, or maintain synchrony across space and among variables (Menzel et al., 2020). Human activities can reduce or disrupt synchrony (Song et al., 2021), such as through irrigation practices designed to maximize crop yields (Kukal & Irmak, 2019; Troy et al., 2015) or through habitat fragmentation (Trenham et al., 2003; Zytynska, 2019). Human activities can also increase or enhance synchrony through biotic homogenization (Olden & Rooney, 2006), seen in large-scale intensive farming in monocultural fields (Benton et al., 2003), and the replacement of native species by exotics during urbanization (McKinney & Lockwood, 1999). Such biotic homogenization can synchronize, for example, plant phenology and pest dynamics (Walter et al., 2020). These local-scale studies suggest complex human modification of ecological synchrony beyond climate change.

In this review, we address the understudied human modification of changing ecological synchrony. We review existing evidence and understandings on the impacts of anthropogenic activities, interacting with climate change, on the spatiotemporal synchrony in ecological systems on different levels of organization (Fig. 2). Despite numerous local evidence, we have a limited consensus and generalized understanding on how ecological synchrony is shaped by management in human-dominated landscapes at regional to continental scales. With ever-increasing data-collection efforts, including observatory networks, remote sensing, and citizen and community science projects, it has become increasingly feasible to systematically investigate the influence of human activities on ecological synchrony. Here, we demonstrate with four case studies from a range of study systems (Fig. 2, Table 1), quantifying the effects of (A) irrigation on ecosystem-level synchrony in croplands, (B) human footprint on community-level synchrony of bird species, (C) managed harvesting on meta-population-level spatial synchrony in marine populations, and (D) temperature on population-level synchrony of mosquito emergence. With these case studies, we demonstrate ways to leverage big data on large spatial scales across levels of organization, thereby enhancing our understanding of human modification of ecological synchrony from a macrosystems perspective.

**Table 1.** Summary of case studies on four levels of organization.

| **Case study** | **A** | **B** | **C** | **D** |
| --- | --- | --- | --- | --- |
| **Level of organization** | Ecosystem | Community | Meta-population | Population |
| **Variables with possible synchrony** | Plant productivity and precipitation | Abundance of birds of different species | Abundance of marine organisms among sites | Emergence of individual mosquitoes |
| **Measure of synchrony** | Correlation between annual summaries | Mean of pairwise correlation coefficients | Decay of correlation over distance | Spread of the timing of events |
| **Human modification** | Irrigation | Human footprint index | Managed harvesting | Possibly urbanization |
| **Data sources** | Long-Term Agroecosystem Research (LTAR) Network, MODIS, Daymet | Breeding Bird Survey (BBS) | Northeast Fisheries Science Center (NEFSC) fall bottom trawl survey | National Ecological Observatory Network (NEON) |
| **Related definition of ecological synchrony** | Coherent temporal changes in biotic and abiotic characteristics | Correlated demographic fluctuations across species (within trophic level) at a single location | Correlated changes in the abundance of geographically disjunct populations | The tendency of individuals to carry out some part of the reproductive cycle at the same time as other members of the population |
| **Previous studies using this definition** | (Özkan et al., 2016; Song et al., 2023) | (Lahoz-Monfort et al., 2013; Royama, 1992; Youngflesh, Li, et al., 2021) | (Bjørnstad et al., 1999; Koenig & Liebhold, 2016) | (Ims, 1990; Youngflesh et al., 2018) |

### Box 1.

Synchrony and asynchrony

Ecological synchrony is traditionally defined as the condition when one or more ecological processes within a system have high concordance or consistent lagged behaviour (Seybold et al., 2021). Conversely, asynchrony is the lack of concordance in fluctuations of ecological processes (Seybold et al., 2021). To clarify, asynchrony usually refers to the lack of a relationship, but not the negative direction of a relationship. For example, the time series of predator and prey population sizes can be exactly out of phase but still in synchrony with each other. In practice, it is often hard to distinguish out-of-phase synchrony, the lack of synchrony, and undetectable synchrony with noisy data (Gouhier & Guichard, 2014).

## 2. Human modifying synchrony across levels of organization

### 2.1. Ecosystem level

Given the profound ecosystem consequences such as nutrient cycling and productivity, examination of the coordination of ecosystem-level processes is warranted (Beard et al., 2019). A unifying framework of ecological synchrony is useful to synthesize theory and data in disciplines such as ecology, biogeochemistry, and hydrology at the ecosystem level (Seybold et al., 2021). Healthy ecosystem functioning requires synchrony between some processes, and asynchrony in some others. In aquatic biogeochemistry, solute concentrations follow synchronous patterns among streams in a watershed, maintaining spatially stable water chemistry (B. W. Abbott et al., 2018). In ecology, loss of asynchrony in biotic processes can destabilize ecosystem functioning (Wang et al., 2021). Such change can be caused by reduced species turnover across space, also referred to as “loss of β diversity” or “biotic homogenization” (Smart et al., 2006). For example, biotic homogenization was shown to synchronize the productivity of grassland communities in biodiversity experiments (Wang et al., 2021). Instead of showing either high degrees of synchrony or asynchrony, ecosystem-level synchrony often follows complex patterns such as synchrony with characteristic lags, spatial synchrony that decays with distance, or changing degree of synchrony (Seybold et al., 2021).

Synchrony between abiotic variables has been examined in several local case studies, with detectable modification by land use. For example, nitrate concentration and stream discharge were generally in-phase in forested and agricultural landscapes, but became more out-of-phase in urbanized landscapes (Van Meter et al., 2020). In another example, human-impacted watersheds had more asynchronous transport of NO ^−^ and dissolved organic carbon across a river network, possibly leading to reduced denitrification reactions by microorganisms (Wymore et al., 2021). Climate change and human modifications interact to influence ecosystem-level synchrony, often altering broad-scale top-down and local-scale bottom-up drivers of synchrony, respectively. For example, dual stressors of increased precipitation and urbanization make stream parameters more spatially synchronized in a watershed, potentially resulting in highly degraded water quality (Vogt et al., 2016). In a terrestrial case study, while climate oscillations naturally induce fire synchrony on a large scale, local land-use modify fire regimes and induce more asynchronous fires among sites (Yocom Kent et al., 2017).

Synchrony involving biotic processes, such as the relationship between the phenology and the environment, are also likely to be modified by human activities, but the effect is less understood. Vegetation phenology shifts rapidly in response to climate change, but has not kept pace with temperature change in human-dominated landscapes, with a greater lag in more densely populated areas (Song et al., 2021). Climate change and urbanization interact to shift both plant reproductive phenology and frost dates, potentially changing plant frost risks (Park et al., 2023). Animals also shift in breeding phenology under warming temperatures, but are at risk of asynchrony with spring temperature changes (Saino et al., 2011), snowmelt (Lameris et al., 2018), or green-up (Mayor et al., 2017). It is yet to be explored how the synchrony between animals and critical ecosystem processes are shaped by human activities. To mitigate the influence of climate change, humans can manage ecosystems to preserve ecological synchrony, such as redesigning sustainable agroecosystems that synchronize the supply of nutrients from soil biota with the fluctuating nutrient demand of plants (Crews & Peoples, 2005; Fontaine et al., 2024) (Box 2).

#### Box 2.

Social-ecological synchrony

The addition of the human modification axis revealed another direction in studying ecological synchrony that is worth further investigation: how humans synchronize our behavior with the environment. During this period of rapid climate change, it is likely that human activities may be increasingly desynchronized with ecological events. For example, tourists’ visits to the wildflowers at a national park are increasingly mismatched with the advancing flowering time (Breckheimer et al., 2020). Meanwhile, we actively organize ourselves in relation to ecological events through practices, rules, norms, and technologies. For example, plant breeding has been used to improve the synchrony of maturity in plant populations to enable more efficient harvests (Doebley et al., 2006; Pickersgill, 2007). Organic farmers who are less reliant on synthetic herbicides tend to plant crops later so that there is a greater asynchrony of crops and weeds, thus reducing weed-crop competition (Cavigelli et al., 2008). Expanding to a sociotechnical level, humans have designed physical (e.g., grain silos) and financial (e.g., futures markets) tools to reduce asynchrony between the supply and demand of commodity crops. The unifying concept of ecological synchrony can be applied to social-ecological systems. Calls have been made for greater ecological intensification in agroecosystems to take advantage of ecological cycles, thereby reducing anthropogenic inputs and harm to intensively managed systems (Bommarco et al., 2013). Framing these issues through the lens of synchrony could be a useful approach to adapting and mitigating the effects of climate change.

### 2.2. Community level

The abundance and species interactions within ecological communities drives the structural and functional stability of ecosystems, and is therefore a main goal for biodiversity conservation and sustainable management (Blüthgen et al., 2016; Hansen et al., 2013; Raimondo et al., 2004). Community-level synchrony has been discussed both on the inter-and intra-annual time-scales, in the contexts of community stability and phenological mismatch, respectively. On an interannual time scale, researchers examine how fluctuations in population of species are coordinated to inform community stability. On an intra-annual time scale, researchers examine how life cycles of species are coordinated to inform phenological mismatch.

Although abundance of single species tend to fluctuate, the total abundance in a community is stabilized, not only by the number of species, but also by their asynchrony, known as the portfolio effect, insurance hypothesis (Blüthgen et al., 2016), or compensatory dynamics (Valdivia et al., 2012; Viviani et al., 2019). Asynchronous fluctuations are often a sign of stable community, with synchronous community-wide fluctuations possibly representing a risk to stability (Gouhier et al., 2010). Note that asynchrony here refers to the lack of synchrony (independent fluctuations), rather than out-of-phase synchrony (negatively correlated fluctuations) (Gouhier & Guichard, 2014; Houlahan et al., 2007) (Box 1). Asynchrony might take an even more central role than biodiversity in promoting community stability (Valencia et al., 2020). A series of metrics have been developed to quantify community-level synchrony, often by comparing the variance of aggregated abundances of taxa to the summed variances of individual taxa (Gouhier & Guichard, 2014; Gross et al., 2014; Hallett et al., 2016; Lepš et al., 2018; Loreau & de Mazancourt, 2008).

Human modifications have frequently been found to increase synchrony among species and thus threaten community stability. Increasing management intensity increased synchrony in plant and animal communities in grasslands and forests (Blüthgen et al., 2016). Human land use and local water pollutants increased synchrony in aquatic communities in a river basin (Li et al., 2023). Anthropogenic activities measured by nutrient concentrations increased synchrony in riverine bacterial communities (Liu et al., 2021). Human management has the potential to promote asynchrony and community stability, such as through herbivore exclusion and reducing grazing intensity, according to a long-term global analysis (Valencia et al., 2020). However, the effects of many management practices are unclear: fertilization and grazing intensification have both been found to have varying effects on synchrony between studies (Blüthgen et al., 2016; Valencia et al., 2020; Zhang et al., 2016).

Phenological mismatch, uncoordinated shifts in the timing of interacting key biological events, is a longstanding field that is increasingly studied under the synchrony framework (Bartomeus et al., 2013; Song et al., 2023). Establishing a baseline for synchrony between phenological events (i.e., no phenological mismatch) is challenging (Lindén, 2018). Some phenological events, such as flowering and pollinator emergence, are expected to be highly synchronous (Lindén, 2018; Renner & Zohner, 2018); some, such as the life history of parasites and hosts, have evolved superficially maladaptive asynchronous strategies (Lindén, 2018; Singer & Parmesan, 2010).

Despite extensive discussion on phenological mismatch driven by climate change (Kharouba & Wolkovich, 2020; Miller-Rushing et al., 2010; Renner & Zohner, 2018; Visser & Gienapp, 2019), the influence of human modification is yet to be systematically examined. Urban phenology has been shown to be modified by multiple factors, including urban heat island effects (Meng et al., 2020), modified water regimes (Buyantuyev & Wu, 2012), artificial light at night (Zheng et al., 2021), and the introduction of non-native species (Alexander & Levine, 2019). For example, increase in the intensity of urbanization has been found to drive an early spring for plants but not for pollinators along the same urbanization gradient, potentially leading to changes in the structure of plant–pollinator networks (Fisogni et al., 2020). In another example, on-native urban trees were shown to have a delayed spring phenology compared to native species, but the phenology of caterpillars better matches their native hosts; this phenological mismatch explains the significantly lower caterpillar abundance on non-native trees, suggesting risks for invertebrate declines in urban areas (Jensen et al., 2022).

### 2.3. Meta-population level

The population dynamics in a meta-population of a species is often spatially synchronous, widely studied for marine organisms (Bjørnstad et al., 1999; A. Liebhold et al., 2004b). Such spatial synchrony is is driven by multiple mechanisms, including coupling through larval dispersal (Molofsky, 1994; Ranta et al., 1998), the Moran effect (Hansen et al., 2020; Moran, 1953; Ranta et al., 1997), and interactions with synchronized species (Blasius et al., 1999; Cazelles & Boudjema, 2001; Ims, 1990). Meanwhile, factors such as heterogeneity in environmental conditions (Lande et al., 1999; Pecora & Carroll, 2015), variable or nonlinear population dynamics (S. B. Hagen et al., 2008; Saether et al., 2008; Stenseth et al., 1999; Swanson & Johnson, 1999), and chaos (Allen et al., 1993; Becks & Arndt, 2013) might reduce or eliminate synchrony (Rogers & Munch, 2020).

Spatial synchrony is a double-edged sword to the persistence of a meta-population. On the one hand, spatial synchrony is often a sign of well-connected meta-populations, with dispersal between populations decreasing population-level extinction risk through the “rescue effect” (Heino et al., 1997; Luo et al., 2021; Plitzko & Drossel, 2015). On the other hand, perfectly synchronous oscillations increase landscape-level extinction risk and threaten species persistence (K. C. Abbott, 2011; Koenig & Liebhold, 2016; Luo et al., 2021; Plitzko & Drossel, 2015). Species persistence in a meta-population is often sustained by a combination of complex patterns of synchrony such as phase synchrony (phase-locked rhythms) (Blasius et al., 1999; Vasseur & Fox, 2009) and asynchrony (including but not limited to chaos) (Blasius et al., 1999; Heino et al., 1997). Therefore, meta-population level synchrony cannot simply be quantified by a single metric and evaluated by a single threshold. Perturbations to the established spatial synchrony can take place in several dimensions, such as correlation strengths, phase differences, and spatial scales.

Human modifications often interact with climatic factors to alter spatial synchrony. Historical overexploitation, compounded by the loss of sea ice corridors under climate change, reduced the spatial synchrony of an Arctic Svalbard reindeer meta-population, threatening the remaining few critically small populations (Herfindal et al., 2022; Peeters et al., 2020). The populations of roe deer in North Germany went through synchronized fluctuations attributed to an expansion in rapeseed cultivation and the North Atlantic Oscillation (NAO) (R. Hagen et al., 2014). In addition, other human modifications that alter population cycles, such as vaccination against pathogens and species extirpations, have been speculated to alter spatial synchrony (Vasseur & Fox, 2009).

A unique topic in meta-population-level (or population-level) synchrony is pest outbreak, with a focus on risks on the persistence of host tree populations. Human modifications have been found to increase spatial synchrony and induce widespread synchronous pest outbreaks through elevated temperature and drought during climate change (Raffa et al., 2008), management practices that favor homogeneous stands of susceptible hosts (Flower, 2016; Raffa et al., 2008), introduction of invasive pests (Potter & Urquhart, 2017), and even pest management practices in conservation lands (Aukema et al., 2006). Taken together, changes in meta-population-level synchrony driven by human modifications often represent challenges to the persistence of populations and species, including interacting species, although the specific consequences are context-dependent.

### 2.4. Population level

While previous case studies focus on ecological synchrony on large spatiotemporal scales and higher levels of organization (ecosystem, community, meta-population), synchrony between organisms within a population is also common but less studied (Lamontagne & Boutin, 2007). In the field of phenology, researchers have observed the prevalence and importance of population-level synchrony. Key life-cycle events such as reproduction often synchronize within a population because of environmental conditions, interactions between individuals, and selection from predators (Loe et al., 2005; Koenig et al., 2015; Fletcher et al., 2010). While most phenological studies focus on point estimates (e.g., first or mean phenological date), assessment of population-level synchrony involves the distribution of phenological dates (e.g., standard deviation, quantiles) (Bolmgren, 1998; Carter et al., 2018; Carter & Rudolf, 2019). Phenological synchrony is crucial for fitness of offsprings (Loe et al., 2005), high fertilization success (Koenig et al., 2015)sal, and escape from predators (Fletcher et al., 2010). Further, phenological synchrony strongly influences intraspecific competition by changing the population density and relative competitive advantages of early- vs. late-arriving individuals (Carter & Rudolf, 2019).

Established phenological synchrony has been suggested to be disrupted by climate change (Koenig et al., 2015; Loe et al., 2005), further complicated by human modifications. One pronounced example is the gradual breakdown of coral spawning phenology in the Gulf of Eilat, reducing the probability of successful fertilization and threatening the population persistence (Shlesinger & Loya, 2019). The loss of coral phenological synchrony was postulated to be driven by changing sea temperature regime and hormonal (endocrine-disrupting) pollutants. In another case, the emergence of salmonfly, an aquatic insect species, was more synchronized at sites in a human-impacted river and less synchronized at sites in a natural river (Anderson et al., 2019), possibly due to changing water temperature regime altered river channels. Such change in synchrony led to a shortened duration of subsidy to consumers of salmonfly. Interestingly, along the entire rivers, salmonfly emergence was more synchronized in a natural compared to human-impacted river, highlighting that the impact of human modification might be scale-dependent.

Apart from synchrony in phenology, individuals in a population might also have distinct synchrony patterns in behaviors, especially in social animals. Synchrony might appear as collective behaviors, a sign of social cohesion, which can be induced or disrupted by human modification. In areas with more pervasive illegal hunting, impala increased behavioral synchrony in response to higher predation risk (Setsaas et al., 2018). The breathing synchrony of pairs of swimming Guiana dolphins has been noted to be disrupted by aluminum research boats, possibly due to interference with acoustic communication (Actis et al., 2018). Synchrony might also appear as alternating behaviors, a form of coordination and cooperation, which again can be altered by human modification in either direction. Parents of house wrens have the behavior of synchronized nest visits, alternating between parents, more in a rural site compared to a suburban one, likely due to the more fragmented habitat and reduced food availability in suburban areas (Baldan & Ouyang, 2020). Human disturbance measured by tourist accessibility changed the form of synchrony from collective to alternating in the antipredator vigilance of black-necked cranes (Kong et al., 2021). Population-level studies together suggest that human modifications have the potential to alter phenological and behavioral synchrony on the population level, but much is unknown on the prevalence, direction, and magnitude of impacts.

## 3. Quantifying human modification with big data

### 3.1. Case study A (ecosystem level)

The tight coupling between primary production and abiotic factors such as temperature and precipitation has been well-documented in the literature, but usually not interpreted in the context of ecological synchrony. We here present the precipitation-productivity relationship as a key example of ecosystem-level ecological synchrony. In particular, we focus on the synchrony in highly managed agricultural systems such as croplands (Weiss et al., 2020) that exhibit unique phenological responses (Taylor & Browning, 2022). Studies with remote sensing data reveal the possibility that not even the rapid shift in plant productivity and phenology match the pace of recent warming (Huang et al., 2017), particularly in human-dominated landscapes (Song et al., 2021). Understanding the nature and degree of synchrony between environmental drivers and primary production or yield in agricultural systems has ramifications for food security and agricultural sustainability (Browning et al., 2021; Mehrabi & Ramankutty, 2019; Spiegal et al., 2018). Here, we use satellite remote sensing data to demonstrate that a less actively managed agricultural system has greater synchrony between primary production and environmental drivers than a more actively managed system, which could be generalized to a larger scale.

We compared managed agroecosystems from two cropland sites with contrasting water management regimes. The selected sites are within 1.6 km of one another at the LTAR Platte River - High Plains Aquifer network location near Mead, Nebraska, USA. The mean annual precipitation is 779 mm, and the mean annual temperature is 10.2 °C (Fick & Hijmans, 2017), where the mean growing season start and end dates are days 150 and 240, respectively (Browning et al., 2021). Both sites are planted with maize-soybean rotations while one site (US-Ne3 RF) is rainfed, and the other (US-Ne2 IR) is irrigated with a center-pivot system. Irrigation management at US-Ne2 IR is based on output from a hybrid-maize model with field-based measurements of evapotranspiration as input (Bhatti et al., 2020).

Satellite remote sensing data on land surface phenology are highly correlated with carbon fluxes measured from eddy-covariance data (Baldocchi, 2020; Novick et al., 2022) and commonly used as a proxy for primary productivity. Specifically, we used a remotely-sensed enhanced vegetation index (EVI) time series at 500 m (MOD13A1v006) every 16 days (Didan, 2015) between 2000 and 2013. We calculated annual productivity metrics (Fig. 3a) using the “greenbrown” R package (Forkel & Wutzler, 2015) after filling winter gaps (Beck et al., 2006; Forkel et al., 2015), temporal smoothing, and scaling (White et al., 1997). Precipitation data at both sites between 2000 and 2013 were obtained from the 1 km Daymet reanalysis product (Thornton et al., 2016a).

**Figure 3.**
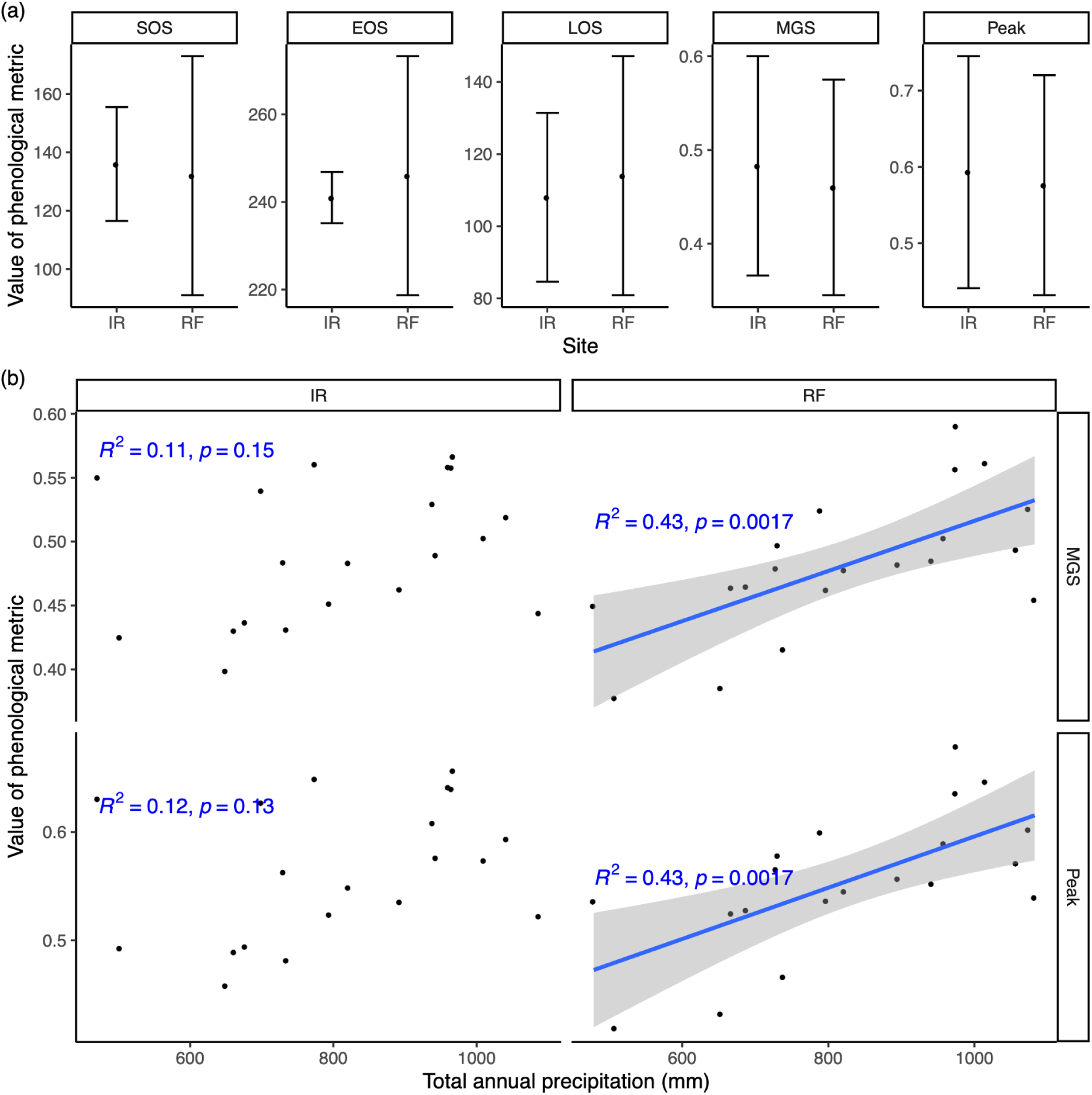
**(a)** Productivity metrics (mean ± standard deviation) including start of season (SOS), end of season (EOS), length of season (LOS), mean growing season EVI (MGS), and annual peak EVI (Peak) for an irrigated cropland (IR) and a rainfed cropland (RF) derived from enhanced vegetation index (EVI) time series from 2000 to 2013. **(b)** Comparison between total annual precipitation and two productivity metrics, Peak and MGS, at IR and RF sites. The coefficient of determination (*R*^2^) and *p*-value are reported for each linear fit. Only fits significant at the *p* < 0.05 level are drawn.

Both sites received similar natural precipitation, while the irrigated site has an additional 187 ± 110 mm of water per growing season (May – September) (Appendix S1: Fig. S1). Phenology metrics indicate less interannual variability in the start of season (SOS), end of season (EOS), and length of season (LOS) at the irrigated site than at the rainfed site (Fig. 3a), suggesting that irrigation decoupled from climatic drivers may homogenize productivity dynamics over the years. We further analyzed the relationships between productivity metrics and precipitation. Notably, peak EVI (Fig. 3b) and mean growing season value (MGS) show significant correlations with precipitation at the rainfed site, but not at the irrigated site (Fig. 3b).

On an ecosystem level, our results indicate that the vegetation productivity in the rainfed cropland is more synchronized with precipitation, compared to the irrigated cropland. More broadly, these results suggest that management interventions like irrigation can potentially change the synchrony between ecosystem processes and climate drivers. It is expected that croplands are managed not to promote ecological synchrony, but to optimize fast growth, high yield, and stress tolerance (Atlin et al., 2017). With rising temperatures and increasingly variable precipitation, agricultural water requirements are projected to increase (Fischer et al., 2007). Common irrigation practices that are based on soil water balance but decoupled from natural precipitation regimes may mitigate loss in crop production during climate change, but at the cost of environmental quality (Kang et al., 2009), such as exacerbating water scarcity (Fujihara et al., 2008) and increasing runoff (Holden & Brereton, 2006). What type of ecological synchrony in human-dominated landscapes we should expect and use as a target for management remains an open question.

### 3.2. Case Study B (community level)

Birds offer an excellent system for studying ecological synchrony, with bird reproductive dynamics often strongly regulated by weather conditions (Shipley et al., 2020), food availability (McKinnon et al., 2012), and interactions with sympatric species (Sanz-Aguilar et al., 2015). These factors have been shown to drive strong within-species population synchrony across space (A. Liebhold et al., 2004a). Studies of population synchrony among species, however, are much less common. Quantifying synchrony at the community level (among species at a given location) has the potential to assist in determining the factors that drive large-scale change and for detecting the effects of human influence.

Using standardized count data on North American birds from 1966 to 2019 obtained as part of the Breeding Bird Survey (BBS) (Robbins et al., 1986), we quantified the degree to which the recorded abundance of passerine bird species fluctuated similarly over time (Fig. 4a). At each survey location (Fig. 4b), we considered only species that were observed in all years. We retained only locations with ten or more species observed in all years and at least ten years of abundance data. In total, data from 3,190 locations were used, with a mean of approximately 19 species at each site (range 10–46) and a mean of approximately 29 years (range 10–54). We detrended the time series of logged abundance to remove any potential effect of similar long-term trends on estimates of synchrony. At each location, we then calculated Pearson correlation coefficients of the logged abundance for each pair of species. We computed the community-level population synchrony for each location as the mean of these pairwise correlation coefficients (Youngflesh, Li, et al., 2021; Gouhier & Guichard, 2014).

**Figure 4.**
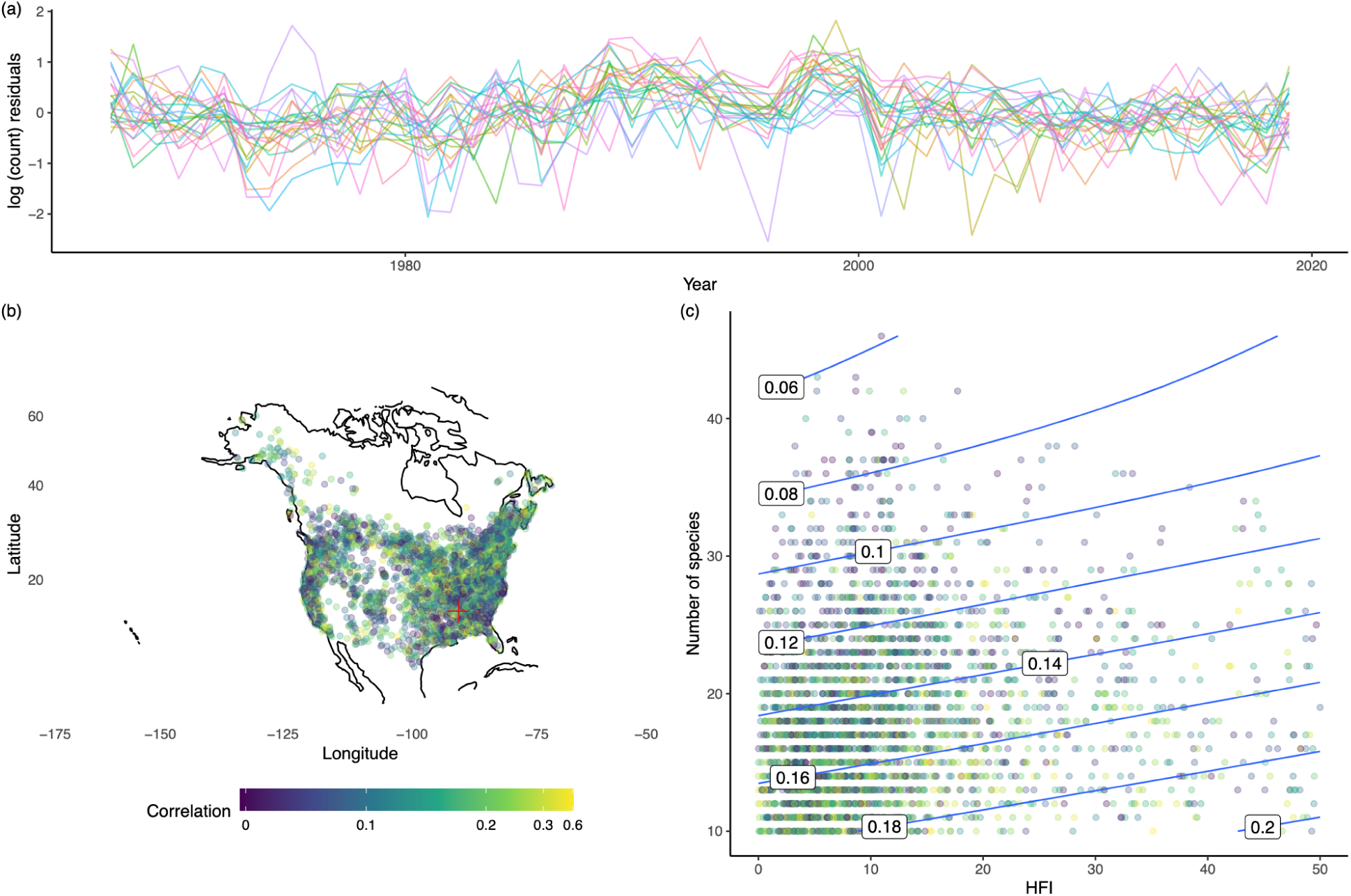
**(a)** Times series of detrended (using a linear regression) of log(abundance) for a community of birds at a random site (St. Florian). Coloured lines each represent a separate species at the site. **(b)** Spatial pattern of community-level demographic synchrony measured in mean pairwise correlation coefficients. The site shown in (a) is highlighted in red. **(c)** Model-predicted community demographic synchrony as a function of the Human Footprint Index (HFI) (a measure of the intensity of human pressure on the landscape) and the number of species in each community. The values on the contour lines correspond to estimates of community demographic synchrony.

We extracted data on the intensity of human impact on these bird communities from the Human Footprint Index (*HFI*), which provides a continuous quantitative estimate (from 0 to 50) of the cumulative pressure on the environment in the year 1993, a midpoint of the study period (Venter et al., 2018b). While the pressure on this environment is likely to have changed over time, we assess synchrony over the entire time period at each location. At each BBS location, we took the average HFI value within a 40 km radius, as the BBS surveys are conducted along a ∼40 km transect. We modeled community demographic synchrony as a function of HFI and the number of species (*N*) in a given community (as more species within a community will make it more likely that community synchrony is near 0 (Loreau & de Mazancourt, 2008)), using a generalized additive model (GAM) (Eqn. 1),

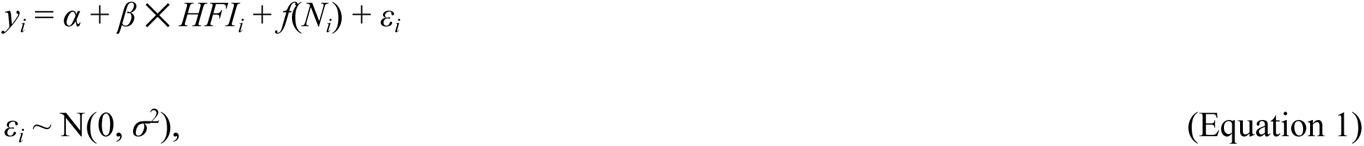

where *α* is the intercept, *β* is the effect of *HFI* on synchrony. *f*(*N*) is a smooth empirical function to characterize the impact of the number of species on synchrony, as we had no prior expectation for the functional form of this relationship. Models were fitted in a Bayesian framework using R package “rstanarm” (Goodrich et al., 2022).

Most locations had positive estimates of community-level population synchrony (0.14 ± 0.10), though estimates varied across space (Fig. 4b). This is in agreement with previous work that has suggested that compensatory dynamics among species are relatively uncommon in ecological communities (Houlahan et al., 2007). The relatively low estimates of community-level synchrony (which could vary from -1 to 1) are also in line with previous work that has suggested that among-species population synchrony may be quite limited (Michel et al., 2016; Youngflesh, Li, et al., 2021), possibly due to niche separation (Sherry, 1979; Youngflesh, Li, et al., 2021) or differential responses to environmental conditions such as climate and land-use change (Michel et al., 2016).

We found a positive association between the intensity of human impact on the landscape and synchrony (posterior mean = 0.0006, 95% CI = [0.0002, 0.0010]) (Fig. 4c). This suggests that either ecological communities in human-dominated landscapes respond more similarly to fluctuations in environmental conditions, or alternatively, experience larger interannual fluctuations in environmental conditions relevant for population processes. Humans might be altering ecological synchrony by altering community composition or inducing converging responses among species in the former case or increasing the magnitude of relevant environmental fluctuations in the latter case. While the precise causal factors of greater synchrony among species in more human-dominated landscapes warrants further exploration, this finding suggests that the persistence and stability of these communities may shift over time, in response to increasing human modification over time (Valencia et al., 2020).

### 3.3. Case study C (meta-population level)

Humans actively manage the populations of marine organisms by changing harvesting efforts based on estimates of current population size and sustainable yields (Methot & Wetzel, 2013), but the effect of such management on spatial synchrony is often overlooked. While traditional optimal harvesting strategies focuses on modeling the dynamics of a single population, recent modeling studies have shown the potential of harvesting to reduce or increase the scale of spatial synchrony, depending on the spatial characteristics of the harvesting strategy (Engen et al., 2018). It is important to avoid the increase of spatial scale, as highly synchronous meta-populations are more likely to undergo unexpected quasi-extinctions (Engen, 2007). We here quantify the impact of managed harvesting on the scale of spatial synchrony with long-term empirical data.

We used data from the NEFSC Fall Bottom Trawl Survey (Azarovitz, 1981) from 1973 to 2016 to examine the spatial synchrony of managed and unmanaged species (Fig. 5a). The standardized NEFSC Fall Bottom Trawl Survey covers a large area from Cape Hatteras, NC, to Nova Scotia, Canada (Fig. 5b), intending to determine the seasonal distribution, relative abundance, and biodiversity of fish and invertebrate species found on the continental shelf in the fall (September to November) in the northeastern United States. We selected 16 species that are abundant in the region and estimated their fall abundance with log-transformed catch per unit effort (CPUE). For each species, we removed sampling sites with zero CPUE in half of the time or more in order to focus on sites with reasonable population sizes.

**Figure 5.**
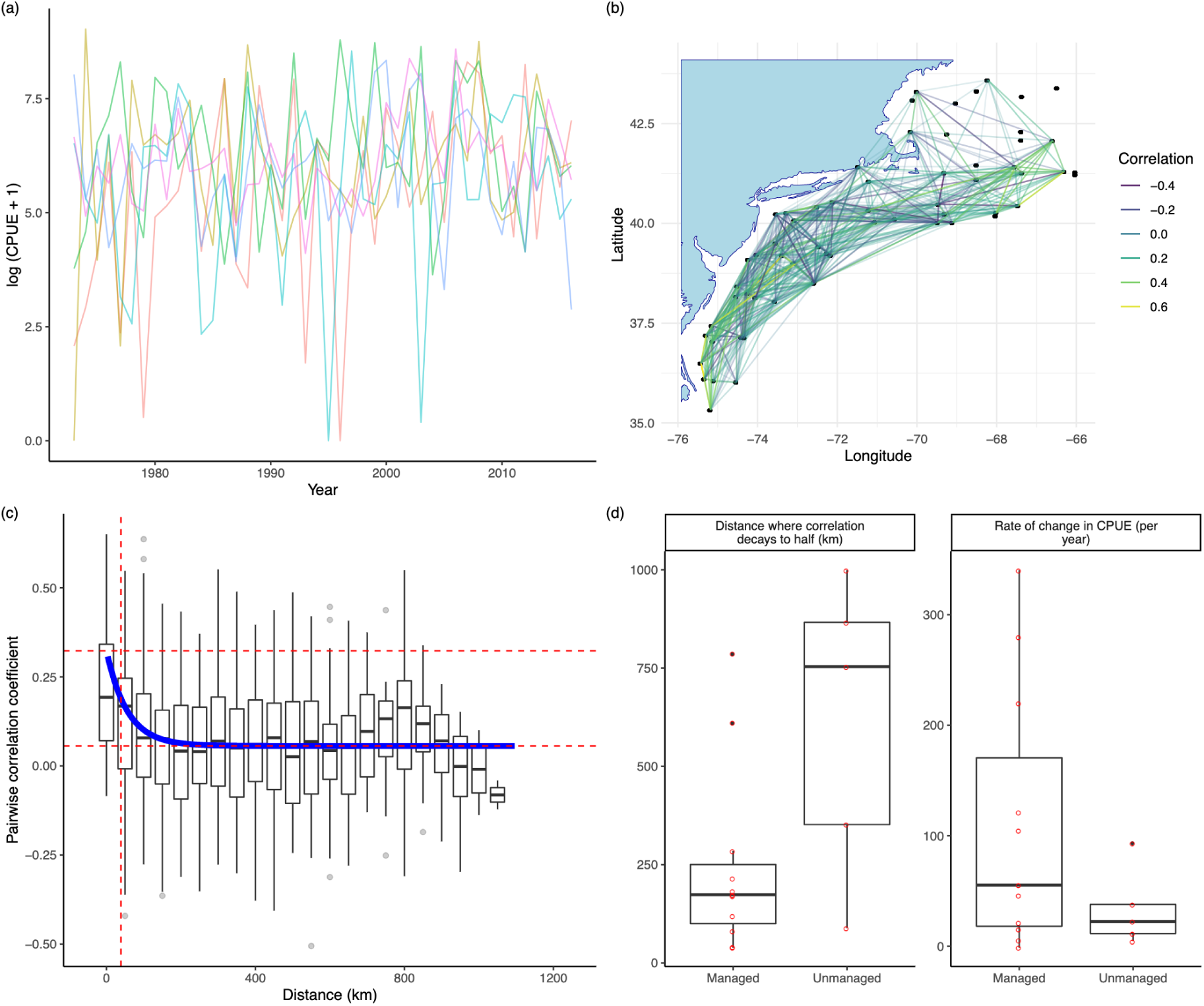
**(a)** Time series of log-transformed abundance at different sites for a marine species, *Loligo pealeii* (longfin squid) as an example of all 16 species. Each colour shows data from a site. Four sites with the highest total abundance are visualised. **(b)** Map of sampling sites (strata) in the NEFSC bottom trawl survey. Colours show pairwise Pearson correlation coefficients in population dynamics between some neighbouring sites. **(c)** Decay of correlation with distance. Blue line shows the fitted exponential decay curve. Red dashed lines show the maximum correlation at a short distance, the minimum correlation at a long distance, and the distance where correlation decays to half. **(d)** Boxplots comparing managed and unmanaged species by distance where correlation decays to half (km) and rate of change in total abundance (CPUE per year).

Similar to case study B, we estimated pairwise temporal correlation in detrended log abundance with Pearson correlation coefficients between sites (Figs. 5b&c). As correlation typically decays with increasing distance, we fitted exponential models to estimate the distance where correlation decays to half (Fig. 5c) to characterize the scale of spatial synchrony (see Appendix S1: Fig. S2 for comparison of other metrics for spatial correlation). Finally, we compared the scale of synchrony between species with and without active management (Fig. 5d). In comparison, we also compared the rate of change in abundance by performing a linear regression of total abundance against year (Fig. 5d).

The mean correlations in managed and unmanaged species were 0.16 ± 0.10 and 0.09 ± 0.05, respectively. Although the mean correlation was significantly higher in managed species (*p* < 0.05), this difference cannot be easily interpreted because of the different geographical distribution of species. The distance at which correlation decayed to half was higher in unmanaged species, 612 ± 380 km, compared to managed species, 246 ± 239 km (*p* < 0.05) (Fig. 5d). This pattern can be compared with that in the rate of change in abundance, where managed species generally have faster increases in abundance (*p* < 0.05) (Fig. 5d). The results suggest that active management of marine population size might not only promote increases in total abundance, but also change the pattern of spatial synchrony. In particular, synchrony may be induced on a smaller spatial scale.

It is encouraging that spatially heterogeneous harvesting did not increase the spatial scale of meta-population synchrony in the area, thus not increasing the risk of quasi-extinction. Even though it is unclear if the maintenance of spatial asynchrony has been an objective in managed harvesting in the area, the result is understandable as harvesting quota is often managed within states or in even smaller units. Given the important implications on the persistence of meta-populations (Earn et al., 1998; Harrison et al., 2020; Schindler et al., 2010), spatial synchrony might be explicitly incorporated into future management considerations.

### 3.4. Case Study D (population level)

We here study a case of mosquito emergence synchrony in nature, showing a viable method that can be used to investigate human influence in the future. Consistent mosquito sampling effort offers an opportunity to study intra-annual synchrony in mobile species populations. Mosquitoes (Diptera: Culicidae) are a diverse family of insects that are aquatic in larval and pupal stages and flying adults. They are both an important food source for other organisms and a common vector of pathogens and parasites. Despite studies on the seasonality of mosquito emergence (Packer & Corbet, 1989), synchronized behavior on the population level is rarely explored. Here, we use the National Ecological Observatory Network’s (NEON’s) continental-scale high-temporal resolution adult mosquito sampling data to examine the ecological synchrony of adult activity within species. While NEON sites are generally located in protected areas with minimal human modification, here we explore environmental conditions that influence synchrony, with application to future research on the role of human activities on mosquito synchrony.

We selected eight NEON sites in the contiguous US with consistent mosquito sampling efforts (Fig. 6A). All sites have low human impacts based on the most recent (2009) Human Footprint Indices (Venter et al., 2018b). NEON mosquito data (DP1.10043.001) (NEON, 2021) for the selected sites during 2016-2020 (Hoekman et al., 2016) were trapping data aggregated to weekly values. We estimated daily abundance as the average abundance per trap hour and processing effort for each sampling event to account for differences in the numbers of hours of mosquito trapping and the proportion of the collected sample processed. Given the gaps and observational noise in data, we modeled seasonal population dynamics for each species by location by year using Bayesian GAMs with the “rstanarm” package in R (Goodrich et al., 2022) (Appendix S1: Text S1).

**Figure 6.**
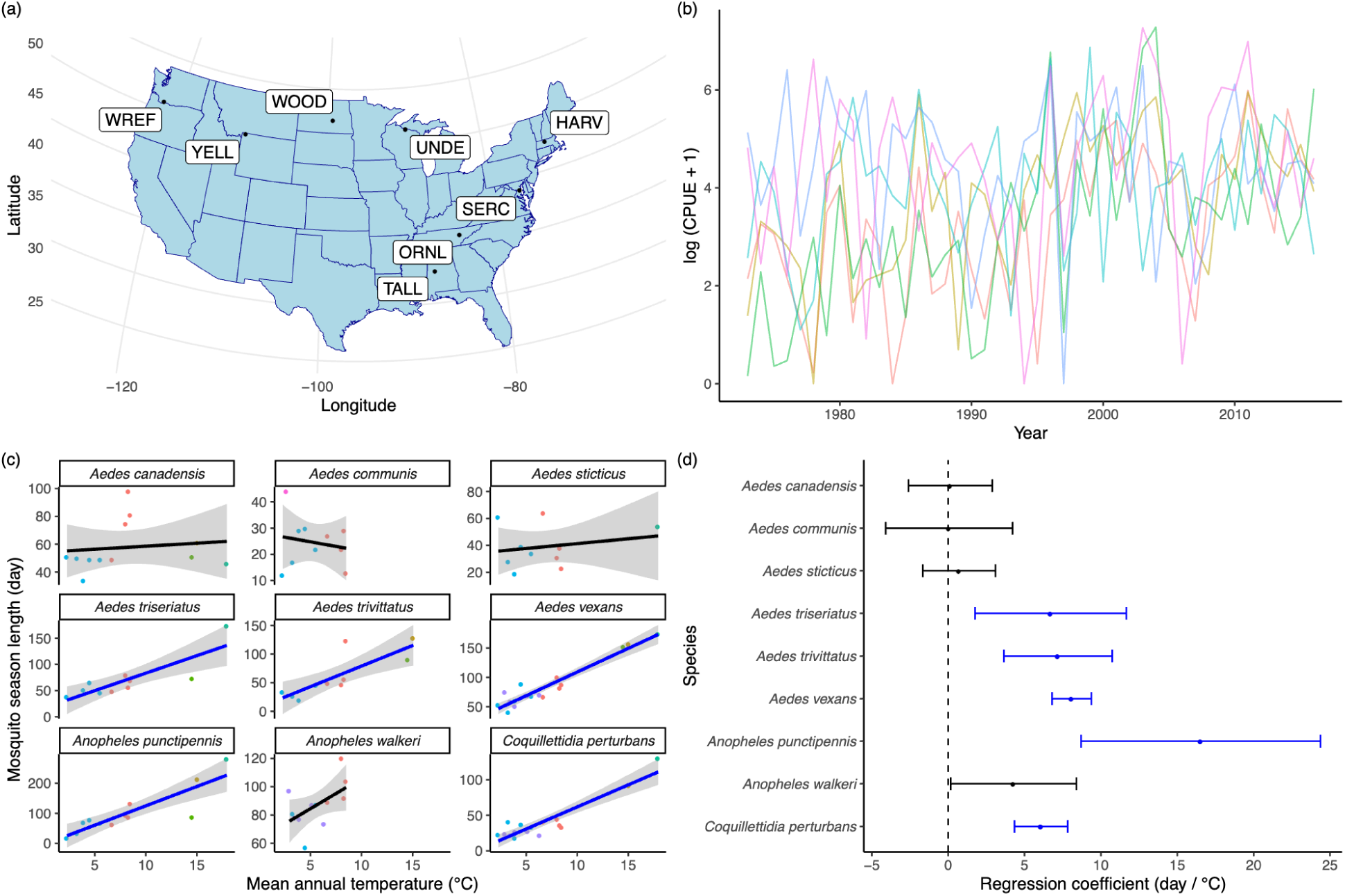
**(a)** Map of NEON sites with adult mosquito sampling data. **(b)** Smoothed time series of the cumulated abundance of *Aedes trivittatus* at Harvard Forest (HARV) site as an example. Shaded areas show the mosquito season marked by the population reaching 10% and 90% of the cumulate abundance at that site and year. **(c)** Correlation between mosquito season length (day) and mean annual temperature (°C). Regression lines with standard errors are shown for each species. Each data point is a site-year, with colours showing different sites. **(d)** Regression coefficients of linear mixed effect models describing the climate-synchrony relationship for each species. Error bars show 95% confidence intervals. Significant effects are shown in blue.

After extracting the fitted seasonal population trends, we quantified the population-level synchrony of adult emergence as the length of mosquito season, i.e., the number of days between the population reaching 10% and 90% cumulated species-specific abundance in the particular site and year. This metric quantifies how synchronized the emergence events are in time: lower season length indicates more synchronized emergence events, whereas higher season length indicates more varied emergence timing among individuals, resulting in a longer duration of species emergence. This metric is commonly used in characterizing reproductive synchrony (Thel et al., 2022). For each species, we assessed how season length correlates with mean annual temperature retrieved from Daymet (Thornton et al., 2016a) by conducting random intercept only linear mixed effect models with the site as random effects using the “nlme” R package (Pinheiro et al., 2017). We focused on species with sample sizes (site ✕ year) greater than 10.

We found that intra-annual synchrony measured by mosquito season length varied among species, year, and site (Fig. 6b). For five out of nine species, there were significant positive correlations (*p* < 0.05) between mean annual temperature (MAT) and mosquito season length, while accounting for the random effect of sites (Figs. 6c&d). The positive correlations are not only the result of climatic differences among sites but also from inter-annual variations between warmer and colder years (Fig. 6c). The result suggests that mosquito species may be less synchronized in warmer years and at warmer sites, leading to more prolonged mosquito seasons. Such relationships between synchrony and temperature, although inferred from data from protected areas, may have implications on the influence of human modifications such as deforestation and urban heat islands.

Exploring the role of human modifications in the population-level synchrony of mosquitoes may have important public health implications, as the degree of synchrony in mosquito emergence may influence the strength and duration of vector disease outbreaks (Yang et al., 2009). Human activities potentially alter the timing of mosquito emergence by urban heat island effects (LaDeau et al., 2015), vector control (Bisanzio et al., 2011), and chemical use (Rochlin et al., 2016). However, gaps in data availability prevent us from analyzing such effects. NEON mosquito sampling is done in relatively less disturbed sites, without an urbanization gradient while data from disease vector control agencies, such as the timing, species, and efforts are not well documented. Although we are not able to test the human modifications to mosquito synchrony, we propose that future mosquito sampling done along urbanization gradients, with both vectors and non-vectors, interpreted in the context of disease vector control, may help us understand this question.

## 4. Synthesis and future directions

In a changing world, future studies on ecological synchrony should explicitly address the role of not only climate change but also aspects of human modifications. On the one hand, the changes of synchrony during climate change may be exacerbated by anthropogenic disturbance; on the other hand, humans may actively influence synchrony to mitigate the negative consequences of climate change. Reviewing the literature on ecological synchrony, we identified multiple streams of evidence on the impacts of human modification, leading to different responses in ecological synchrony in human-dominated landscapes compared to natural landscapes. With four case studies from distinct systems, we demonstrated methods to identify footprints of human modifications on ecological synchrony with big data on large scales across levels of ecological organization. Such quantitative analyses have revealed various types of anthropogenic effects on ecological synchrony, such as disrupted correlation between ecosystem processes, loss of asynchrony between species in a community, and changes in scales of spatial synchrony in meta-populations.

Despite ecological synchrony being a highly complex topic, we identified several insights recurring throughout our literature review and case studies.

1. Human modification has the potential to alter ecological synchrony across scales, levels of organization, and study systems, often with ecological and societal consequences.
2. Direction, magnitude, and consequences of human-modified synchrony are context-dependent.
3. Climate change and human modifications interact to influence (a)synchrony, often with climate change acting as broad-scale, top-down drivers, and human modifications as local-scale, bottom-up drivers.
4. Studying anthropogenic ecological changes through the lens of ecological synchrony allows us to draw connections across levels of organization with generalizable theory (e.g., complementarity, mismatch) and statistical methods.
5. Big data allows human modification of synchrony to be studied on large spatial scales and across levels of organization, enhancing our understanding from a macrosystems perspective.

There remain key unanswered questions in our understandings of ecological synchrony. The collection and publication of big data, from observatory networks, remote sensing, and citizen and community science projects, provides opportunities to answer these questions on large scales and cross-discipline collaborations. We here outline a few questions that could potentially be addressed with big data, with a focus on data from observatory networks. Examples of such observatory networks include the National Ecological Observatory Network (NEON) (Loescher et al., 2016), the Long-term Agroecosystem Research Network (LTAR) (Spiegal et al., 2018), the Long-term Ecological Research Network (LTER) (Vanderbilt & Gaiser, 2017), and the Critical Zone Observatories (CZO) (Guo & Lin, 2016). We also identify challenges and priorities for observatory networks.

1. Do human modification of synchrony scale from local to global scales? Despite extensive local case studies, continental to global scale consensus on the impact of human modification on synchrony is rare, leaving uncertainty in its large-scale patterns and consequences. To systematically and quantitatively understand the impacts of human modification on synchrony requires observatory networks to collect data at broad spatial extents across gradients of human modification. Although existing observatory networks have large spatial coverage, they are often spatially sparse and lack human modification gradients within networks (e.g., NEON and LTAR), calling for in-depth collaboration between and beyond networks.
2. Can we predict the impacts of human modification on synchrony? Moving beyond descriptive and correlational studies that give highly context-dependent insights, it is necessary to make predictions of ecological synchrony in a changing future for mitigation and adaptation. Predictions might be achieved through advancement in data and models that explicitly account for the spatiotemporal patterns of human modification drivers, climate change drivers, and focal ecological processes. A challenge exists in the curation of human modification data in observatory networks, asking for better documentation of the time, location, type of disturbances and management practices, as well as connections to natural ecological data.
3. What is the historical baseline of synchrony, prior to extensive climate change and human modifications? The baseline synchrony is highly context-dependent. For example, plant masting have long been synchronized within populations (Pesendorfer et al., 2021). Phenology of interacting species have evolved synchrony for mutualistic relationships (e.g., plant-pollinator) or asynchrony for antagonistic relationships (e.g., predator-prey) (Lindén, 2018; Singer & Parmesan, 2010). Further, the abundance of marine populations might be expected to be weakly synchronized over space due to natural environmental heterogeneity (Rogers & Munch, 2020). The timespan of data from observatory networks poses a challenge to study historical baselines, but museum collections and automated collection of imagery and audio records present new opportunities to examine the changes in synchrony over time.
4. What should be the target(s) of sustainable management of synchrony? Optimal ecological synchrony for key ecosystem services is different and ever-changing. A certain level of synchrony is necessary to maintain the genetic diversity in a small and fragmented meta-population (Peeters et al., 2020). High synchrony in pests and disease vectors needs to be addressed as it threatens natural biological control. The use of N fertilizers desynchronized the supply and demand of nitrogen to increase yield for provisioning, but management practices that promote their synchrony are now encouraged to restore ecosystem regulating for soil, groundwater, and air (Crews & Peoples, 2004). The goal-setting for managing ecological synchrony requires inclusive social science studies possibly in collaboration with observatory networks.
5. How does changes in synchrony cascade through levels of organization? Although changes in synchrony have been studied across individual levels of organization, there has not been extensive research on the “cascade of ecological synchrony” through cross-scale interaction and cross-scale emergence (Heffernan et al., 2014). For example, changes in the emergence between pest individuals on a population level might scale up to changes in the interaction between pest and host plant phenology on the community level (Grulke, 2011), and further to the pace of nutrient input and uptake on the ecosystem level (Grüning et al., 2017). Strong collaboration among researchers is needed to bridge subdisciplines in ecology to examine the cascade of ecological synchrony from a macrosystems perspective.

## Acknowledgements

This research was initiated during a virtual workshop “Complex Landscapes at Scale: Integrating our Understanding of Managed and Unmanaged Lands at Regional to Continental Scales” (May 27 – June 25, 2021), funded by the National Science Foundation (NSF Award 2044293 to Michigan State University). Y. Song was supported by the UCSC Hammett Fellowship and the Eric and Wendy Schmidt AI in Science Postdoctoral Fellowship, a Schmidt Sciences program. D. M. Browning was supported by USDA CRIS fund 3050-11210-009-000. B. Zuckerberg was supported by NASA Awards 80NSSC19K0180 and 80NSSC21K1348 and NSF Award 1926428. K. M. Dahlin was supported by NSF Awards 1702379 and 2044818 and the USDA NIFA Hatch project 1025001. A. Bybee-Finley was supported by NIFA Grant 2019-67019-29468. C. Youngflesh was supported by NSF Award 2033263 and a Presidential Postdoctoral Fellowship from Michigan State University. K. Zhu was supported by the UCSC Committee on Research Faculty Research Grant and NSF Awards 1926438, 2306198 (CAREER). This research was a contribution from the Long-Term Agroecosystem Research (LTAR) network. LTAR is supported by the United States Department of Agriculture. Case study D was conducted as part of the “Forecasting Mosquito Phenology in a Shifting Climate: Synthesizing Continental-scale Monitoring Data” Working Group supported by the John Wesley Powell Center for Analysis and Synthesis, funded by the U.S. Geological Survey. We thank Dr. Tanya Rogers for her help in retrieving and pre-processing NEFSC Fall Bottom Trawl Survey data.

## Author contributions

All authors collectively conceived and designed the study. Y. Song developed and drafted the theoretical framework. Y. Song, M. Barnes, D. M. Browning, K. M. Dahlin, S. B. Munch, C. Youngflesh, and K. Zhu conducted data analyses for case studies and drafted corresponding sections. Y. Song, A. Bybee-Finley, G. E. Ponce-Campos, and B. Zuckerberg wrote parts of the introduction and synthesis. The manuscript received substantial input and review from all authors.

## Data availability statement

We used published and publicly available raw data in all case studies. MODIS Enhanced vegetation index data MOD13A1 v006 (Didan, 2015) are available from NASA EOSDIS Land Processes Distributed Active Archive Center (LPDAAC): https://doi.org/10.5067/MODIS/MOD13A1.006. Daymet data version 3 (Thornton et al., 2016b) are available from ORNL Distributed Active Archive Center (ORNLDAAC): https://doi.org/10.3334/ORNLDAAC/1328, here downloaded using the “daymetr” R package (Hufkens et al., 2018). Breeding Bird Survey data (Robbins et al., 1986) are available from USGS ScienceBase: https://doi.org/10.5066/P97WAZE5, here downloaded using the “bbsBayes” R package (Edwards & Smith, 2021). Human footprint data (Venter et al., 2018a) were downloaded from NASA Socioeconomic Data and Applications Center (SEDAC) at https://doi.org/10.7927/h46t0jq4. NEFSC Fall Bottom Trawl Survey data (Azarovitz, 1981) are available from Data.gov: https://catalog.data.gov/dataset/fall-bottom-trawl-survey. Mosquito sampling data (NEON, 2021) are available from NEON (National Ecological Observatory Network) data portal: https://doi.org/10.48443/c7h7-q918.

