## Supplementary Information for "Ecological synchrony in human-modified landscapes under a changing climate"

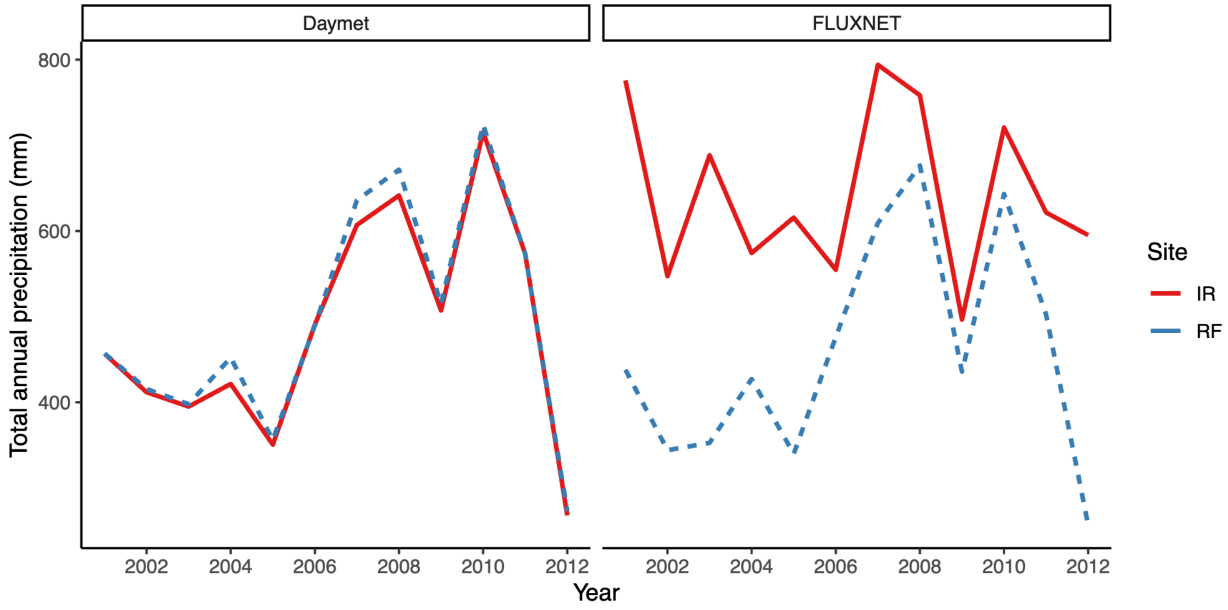


**Figure S1.** Comparison of total annual precipitation (mm) between an irrigated cropland (IR) and a rainfed cropland (RF) at the LTAR Platte River - High Plains Aquifer network location, as calculated from **(a)** reported FLUXNET tower-measured precipitation and **(b)** Daymet gridded precipitation (1 km resolution).

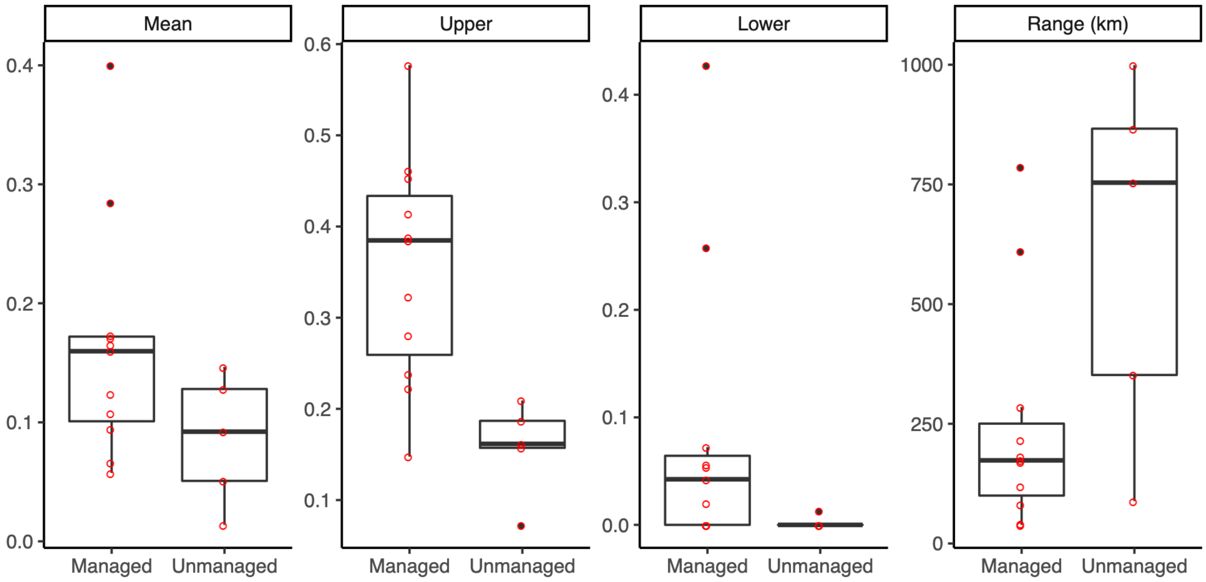


**Figure S2.** Comparing the spatial correlation of managed and unmanaged marine species on the continental shelf of northeastern United States from 1973 to 2016 in terms of the mean correlation among all pairs of sites (Mean), the maximum correlation at a short distance (Upper), the minimum correlation at a long distance (Lower), and the distance where correlation decays to half (Range) (km).

**Text S1.** Preprocessing mosquito population dynamics data.

We first modelled population dynamics for each species by location by year using bayesian general additive models (GAMs) with the "rstanarm" package in R (Goodrich et al., 2022). As different mosquito species have different life histories and may respond to different environmental cues, we did not expect them to have a simple unimodal relationship with seasonality and instead exhibit a wide range of population dynamics including variation in skewness and number of seasonal peaks. Using GAMS provides the flexibility to account for these nonlinear dynamics while allowing us to account for variation in sampling effort (i.e., variation in trap hours and proportion of sample processed). We fitted the daily population abundance for each species ✕ site ✕ year combination using a cyclic cubic spline smoother for day of year with an offset that accounts for the proportion of sampled processed and the total amount of trap-hours. Each model was run using four chains and 3000 iterations, model fit was evaluated by assessing convergence and the rhat factor. After fitting the model, we extracted the median best fit line from the posterior distribution of the predictor variables, used in the following analyses of synchrony.
